# Molecular mechanisms of immune evasion by host protein glycosylation of a bacterial immunogen used in nucleic acid vaccines

**DOI:** 10.1101/2025.05.29.656815

**Authors:** M.S. Cinar, T.M. Adams, Z. Nawaz, E.S. Demir, M.E. Demirturk, A.P. Keelaghan, S.M. Nazaar, B.R. Roberts, A. Ozdilek, F.Y. Avci

## Abstract

Nucleic acid vaccines (DNA and mRNA) induce immunity by driving in situ antigen expression in host cells. For non-viral pathogens, however, host expression can impose post-translational modifications absent from the native microbial antigen. Tuberculosis (TB) remains a leading cause of infectious mortality, and nucleic acid vaccines targeting the *Mycobacterium tuberculosis* antigen 85 (Ag85) complex did not confer protective efficacy in clinical trials. We hypothesized that host-derived N-glycosylation of Ag85 immunogens expressed in mammalian cells compromises immune recognition. Here, we define structural, biochemical and immunological mechanisms by which host-imposed N-glycosylation remodels a bacterial antigen expressed in mammalian cells. We show that Ag85B expressed in human Expi293 cells is microheterogeneously N-glycosylated at four canonical sequons (N52, N224, N234, N280) with predominantly complex, highly fucosylated, and frequently sialylated glycans. Molecular dynamics simulations indicate that these glycans occupy substantial conformational space and reduce solvent and antibody-accessible surface area, occluding multiple established B-cell and T-cell epitope regions. Consistent with glycan-mediated shielding, mammalian-expressed Ag85B shows markedly reduced binding to an Ag85-complex monoclonal antibody by competitive ELISA and biolayer interferometry, and sialylated glycans enable Siglec-9 binding that is abrogated by sialidase treatment. Together, these findings define the structural and biochemical mechanisms by which host glycosylation can remodel bacterial vaccine antigens, supporting glycosylation-aware immunogen engineering as a design principle for nucleic acid vaccines targeting non-viral pathogens.

## Introduction

Nucleic acid vaccines, including DNA, recombinant viral vector, and mRNA platforms, represent a next-generation immunization strategy that diverges from conventional vaccine modalities (1). Instead of delivering whole pathogens or purified protein antigens, these vaccines introduce the genetic sequences encoding pathogen-associated proteins directly into host cells, expressing the target antigens *in situ*. This platform offers significant advantages in manufacturing scalability, flexibility, and rapid adaptability, particularly in response to emerging infectious diseases (2).

While nucleic acid vaccines have long been studied in both human and veterinary contexts, they were not approved for human use until the global urgency of the COVID-19 pandemic catalyzed the clinical deployment of several mRNA-based vaccines (3–6). This landmark success has revitalized interest in using nucleic acid vaccine platforms for prokaryotic and eukaryotic pathogens.

However, an essential immunological distinction emerges when applying nucleic acid vaccine technology to non-viral pathogens. Unlike viral proteins, which are naturally synthesized within host cells during infection, non-viral antigens are not expressed in host cells upon infection. We postulated that employing nucleic acid vaccines against non-viral pathogens would inadvertently render the proteins expressed by host cells structurally and immunologically distinct from their native counterparts expressed by the pathogen. This mismatch primarily arises from the host’s post-translational modifications (PTMs), especially glycosylation for targets entering the secretory pathway. Glycans may obscure or alter immunogenic epitopes and/or engage immune regulatory pathways that suppress effective responses (7, 8). Such host-derived glycans could therefore compromise vaccine efficacy by reducing the quality and magnitude of immune recognition and function.

This concept is particularly relevant for tuberculosis (TB), caused by *Mycobacterium tuberculosis* (*Mtb*), which remains one of the deadliest infectious diseases worldwide (9, 10). With an estimated one-third of the world’s population infected by *Mtb,* over ten million new TB cases, and over one million deaths annually, TB presents an urgent global health challenge exacerbated by the emergence of multidrug- and extensively drug-resistant *Mtb* strains (9, 11). Current treatments are lengthy, expensive, and often not curative, emphasizing the need for more effective vaccines (9, 12, 13). The only approved vaccine, Bacillus Calmette-Guérin (BCG), is an attenuated *Mycobacterium bovis* strain introduced nearly a century ago and confers limited protection (14). It particularly fails to prevent adult pulmonary TB, necessitating the development of protective new-generation vaccines (15–18).

Among the most promising TB vaccine targets employed in multiple vaccine clinical trials are antigen 85 (Ag85) proteins, Ag85A and Ag85B—fibronectin-binding mycolyltransferases critical to *Mtb* cell wall assembly and survival (19). These proteins are abundantly expressed on the outermost surface of *Mtb*, highly immunogenic, and essential for intracellular replication, making them high-priority targets in multiple clinical vaccine trials (20). However, despite their promise, Ag85A- and Ag85B-based nucleic acid vaccines failed to elicit protective immunity in several clinical trials (21–26).

Our prior work revealed that Ag85A produced in mammalian expression systems undergoes N-glycosylation, which significantly dampens its immunogenicity compared to the native *Mtb*-expressed form (27). Building on this foundational work in a mechanistic direction, we dissect how host-imposed N-glycosylation modifies Ag85B, an extensively pursued TB vaccine antigen, by defining the structural and biochemical pathways by which non-native glycans impair epitope accessibility and immune recognition. Using site-resolved glycoproteomics/glycomics, molecular dynamics simulations, and quantitative antibody binding, epitope mapping, and immune interaction assays, we identify how glycan occupancy, microheterogeneity, and terminal sugar composition contribute to antigenic masking and potential immunoregulatory receptor engagement. These findings inform the development of design principles for nucleic acid vaccines against non-viral pathogens, where maintaining native-like antigen structure may require removing vulnerable sequons and/or controlling antigen trafficking through glycosylation pathways.

## Results

### Ag85B is microheterogeneously glycosylated at four canonical N-linked glycosylation sequons when expressed in human Expi293 cells

Mammalian N-glycosylation takes place canonically at N-x-S/T(C) sequons, where “x” may be any amino acid except proline (28). *Mycobacterium tuberculosis* Ag85B (*Mtb* Ag85B) is not N-glycosylated when expressed in its native form; however, the protein sequence predicts that four canonical N-glycosylation sites may be glycosylated if expressed in mammalian hosts **(Fig. 1a)**. To mimic the protein expressed by nucleic acid vaccines targeting Ag85B, we expressed the Ag85B protein (293-F Ag85B) in Expi293 cells, a variant of a human embryonic kidney cell line (HEK293-F), and investigated its potential glycosylation.

**Figure 1:**
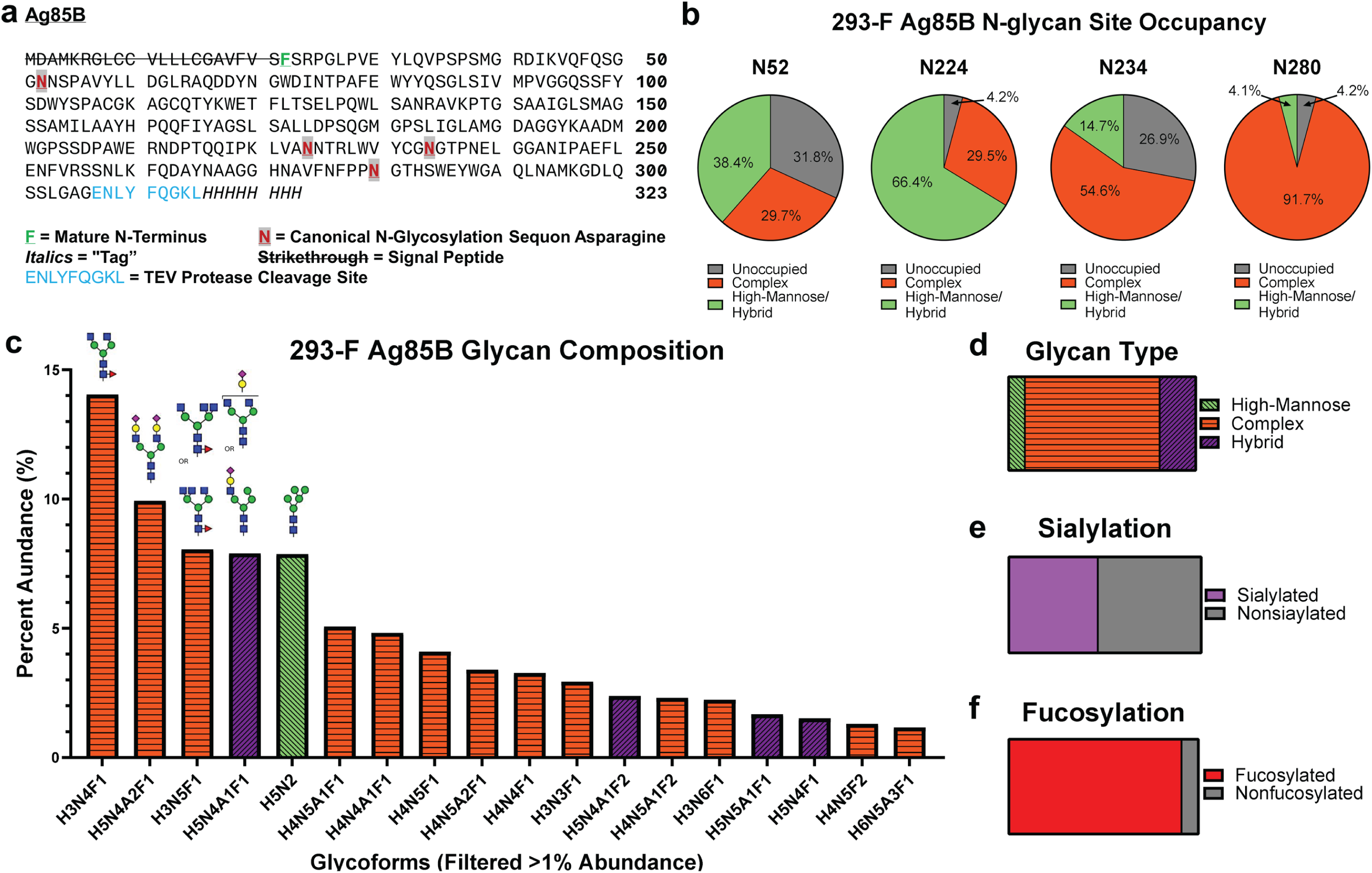
Ag85B is glycosylated at four canonical N-linked glycosylation sequons when expressed in mammalian Expi293 cells. **(a)** Protein sequence of Ag85B. The mature N-terminal is bolded, underlined, and shown in green. The canonical N-linked glycosylation sequon asparagine is bolded in red text, highlighted in gray, and underlined. The TEV protease cleavage site is displayed in blue text, and the 8xHIS tag is italicized. **(b)** Site occupancy of canonical N-linked glycosylation sites (N52, N224, N234, and N280) using EndoH and PNGase F digestion with heavy water, D_2_O. **(c)** Glycomics data showing the percent abundance of released glycans from 293-F Ag85B using PNGase F digestion filtered to show greater than 1% abundance in the total identified glycoforms. Complex structures are represented in dotted orange, hybrid structures are represented in checkered purple, and high-mannose structures are represented in triangle light green. Putative structures shown for the most abundant glycan compositions; note that these compositions are a mixture of isomers. **(d)** Distribution of high-mannose (8.7%) and (72.0%) complex, and (19.3%) hybrid glycoform structures from the total identified glycoforms. **(e)** Distribution of sialylated (46.3%) glycoforms from the total identified glycoforms. **(f)** Distribution of fucosylated (91.1%) glycoforms from the total identified glycoforms.

To assess the N-glycosylation of Expi293-expressed (glycosylated) Ag85B, we first conducted N-glycan site-occupancy analysis using nLC-MS/MS mass spectrometry and an EndoH/PNGase F glycosidase with heavy water (H_2_^18^O) labeling scheme (29). This method simplifies N-glycan analysis by binning N-glycan sites into one of three potential groups: unoccupied (+0 mass shift), high-mannose/hybrid (+203 mass shift), and complex (+3 mass shift). The differentiation between high-mannose/hybrid and complex sites is based on EndoH’s recognition of terminal mannose groups. Our study confirmed N-glycan presence in the four canonical sequons: N52, N224, N234, and N280 **(Fig. 1a)**. Our analysis revealed that N-glycans heavily occupy these four sites, with the lowest occupancy being 68.1% at site N52 (sequon N_52_NS) and the highest occupancies being 95.8% at both sites N224 (sequon N_224_NT) and N280 (sequon N_280_GT) **(Fig. 1b)**. This also aligns with the known preference that NxT sites are favored by oligosaccharyltransferase (OST) over NxS sites (30).

Glycomics analysis of glycosylated Ag85B by LC-MS revealed a heterogeneous population of glycoforms. A total of 75 different glycan compositions were detected, 18 of which are present with greater than 1% of total glycan abundance **(Fig. 1c)**. Additionally, 72.0% of all compositions were identified as complex glycoforms **(Fig. 1d)**. The most abundant composition identified is the complex type N-glycan HexNAc(4)Hex(3)Fuc(1), accounting for about 14% of all glycoforms present. Moreover, several structures with greater than 1% abundance exhibit varying degrees of sialylation and core fucosylation **(Fig. 1c)**. These data are generally consistent with the site occupancy results, with the discrepancies between the two experiments likely due to the hybrid glycan classification scheme used in our glycomics approach.

In addition to the classical roles of N-glycans in contributing to glycoprotein structure and stability, sialylation and fucosylation of N-glycans play roles in immune modulation, including self-recognition, in the human immune system (8, 31, 32), with particular importance placed on the interaction of sialic-acid binding immunoglobulin-like lectins (siglecs) with sialic acids. Furthermore, bacterial species can use terminal sialic acid and fucose moieties to engage in “molecular mimicry” and evade immune responses (8, 33–35). To assess the presence of these immunoregulatory terminal sugars, we analyzed the extent of sialylation and fucosylation of these proteins during post-translational modification. Of all mammalian-expressed Ag85B glycans, 46% are sialylated, and 91% are fucosylated. The prevalence of such moieties on the surface of the mammalian-expressed Ag85B may alter its immune recognition **(Fig. 1e-1f)**.

To better understand the specific glycan profile at each identified sequon, we performed glycomics-informed glycopeptidomics analysis using mass spectrometry (36). A representative MS2 spectrum of the N52 glycan site demonstrates the rich fragmentation generated from our stepped high-energy collisional dissociation (sHCD), which provides functional fragments both of the peptide backbone and of the glycan itself in a single MS2, which are used by the search engine pGlyco3 to identify glycopeptides **(Supplementary Fig. 1)**. N-glycan microheterogeneity (site-specific diversity and distribution (37)) was observed at all sequons. The most abundant glycan at site N52 was the high-mannose structure HexNAc(2)Hex(5) **(Supplementary Fig. 2a)**, which agrees with site occupancy data showing an abundance of terminal mannose structures at that site. Site N224 contrasts this with abundant complex glycosylation and complete fucosylation of non-oligomannose structures **(Supplementary Fig. 2b)**. At site N234, a mixture of oligomannose, hybrid, and complex structures coexist with the more processed glycans appearing to be biantennary fucosylated structures **(Supplementary Fig. 2c)**. Finally, site N280’s glycopeptide analysis data contradicts the site occupancy data with the most common glycoform being the oligomannose HexNAc(2)Hex(5) N-glycan, which commonly forms a bottleneck at MGAT1 processing **(Supplementary Fig. 2d)**. One explanation for this is the higher branching at this site (as evidenced by more HexNAc groups) ‘spreads’ the glycoform distribution across a broad range of potential structures, diluting the signal. Ion suppression may also play a role, mainly because this site differs from the others in that it requires a non-tryptic peptide, which is not ideal for analysis. However, the previously observed patterns of high fucosylation levels and abundant sialylation among the more processed glycans remain consistent.

### Molecular dynamics simulations indicate N-glycosylation decreases the surface accessibility of Ag85B

To better illuminate the significant impact of N-glycosylation on the structure and surface accessibility of Ag85 B, we performed molecular dynamics (MD) simulations. The glycosylated protein (Expi293-expressed Ag85B) was generated using GLYCAM (38, 39) to add site-specific, abundant glycan structures, identified through glycomics-informed glycopeptidomics mass spectrometry data **(Supplementary Fig. 2)**, to a previously published *Mtb* Ag85B X-ray diffraction structure, 1F0N (40). As a control and for comparison, we also simulated the native, nonglycosylated protein structure. We evaluated surface accessibility to water and to larger molecules, such as antibodies, using probes of varying sizes and software tools.

First, solvent-accessible surface areas (SASAs) were assessed using a 1.4 Å probe corresponding to the radius of a water molecule, with CPPTRAJ (41). SASA values were calculated for each frame across the 1-μs production replicates and plotted over time **(Fig. 2a)**. Throughout the MD simulation, the glycosylated protein consistently exhibited a significant reduction in water accessibility across nearly all conformations sampled when compared with the nonglycosylated protein. Although one replicate from each condition showed some overlap, these points correspond to the lowest accessibility values in the nonglycosylated protein and the highest in the glycosylated protein. When re-examined as distributions in violin plots, the reduction in solvent accessibility in the glycosylated protein becomes even more apparent **(Fig. 2b)**. The mean SASA for the nonglycosylated protein was 116 nm². In comparison, the glycosylated protein had an area of 107 nm². Overall, as demonstrated by the mean SASA plots, the two proteins show distinct accessibility profiles **(Fig. 2a-2b)**.

**Figure 2:**
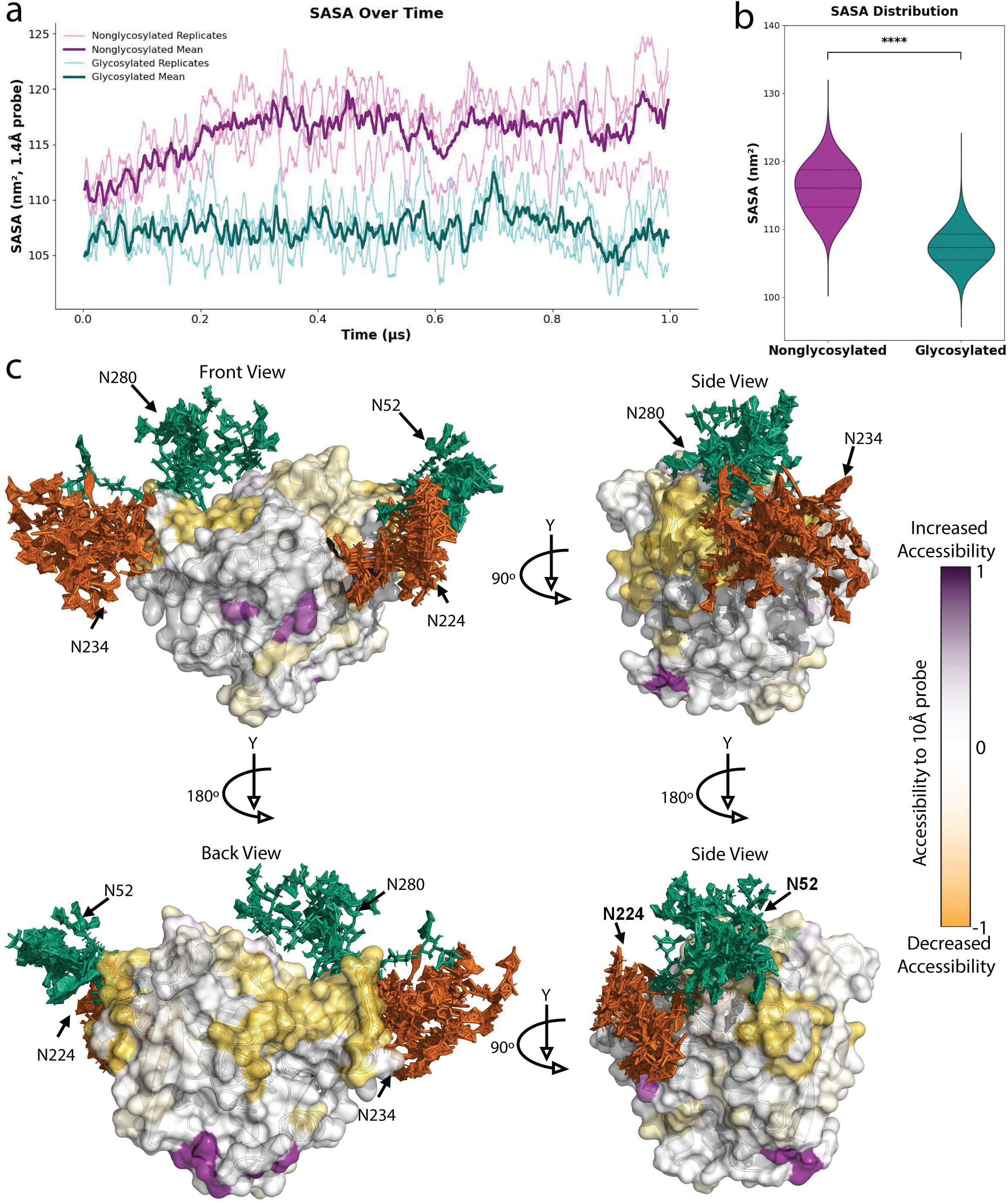
Molecular dynamics simulations indicate the impact of Ag85B N-glycosylation on the protein surface accessibility. **(a)** Water accessibility over three 1 μs-long molecular dynamics simulations of the nonglycosylated and glycosylated Ag85B with a probe size of 1.4 Angstroms. Replicates are displayed in a lighter shade of magenta and teal for the nonglycosylated and glycosylated MD runs, respectively. Mean of all three replicates displayed in a darker shade of magenta and teal for the nonglycosylated and glycosylated MD runs, respectively. **(b)** The water-accessible surface area from (a) is demonstrated as a distribution. ∗∗∗∗*p* < 0.0001 represents a significant difference between groups determined using the Mann-Whitney U test. **(c)** Hierarchical ensemble representation of the glycomics-informed glycopeptidomics site specific abundant glycan structures of Ag85B generated after three 1 μs long molecular dynamics simulations viewed from the front view (top left), back view (bottom left), side view when rotated 90 deg to the left in respect to the front view (top right), and side view when rotated 90 deg to the left in respect to the back view (bottom right). The gradient bar demonstrates the percent change in accessibility to a 10 Angstrom probe, representing the antibody-accessible surface area of the glycosylated Ag85B protein compared to the native, nonglycosylated protein. Residue-specific antibody-accessible surface areas were calculated using NACCESS. Positive/purple represents increased antibody accessibility, and negative/yellow represents decreased antibody accessibility. Complex/hybrid glycan structures are represented in orange. High-mannose structures are represented in green. The protein backbone is represented in white.

Next, to assess antibody-accessible surface areas (ASAs), we used a larger 10 Å probe with NACCESS, which provides residue-specific accessibility calculations (42, 43). To simplify NACCESS analysis, we used hierarchical clustering to reduce the glycosylated protein trajectory to ten representative structural conformations **(Supplementary Fig. 3a-3b)**. Figure 2c illustrates various angles of each glycan’s conformational flexibility across these ten structures, highlighting the vast three-dimensional space that glycans can occupy on the protein surface over time. When assessing residue-specific ASAs, regions of the protein backbone closest to glycans consistently showed decreased accessibility. In contrast, residues distant from glycosylation sites either showed no change or increased accessibility **(Fig. 2c)**. Together, our SASA and ASA results demonstrate that N-glycans occupy substantial surface regions on Ag85B, leading to measurable reductions in solvent and antibody accessibility. These structural alterations may influence epitope exposure or shielding, potentially affecting recognition by humoral immune components. This may result in the shielding of vulnerable antigenic targets and the exposure of non-productive epitopes.

**Figure 3:**
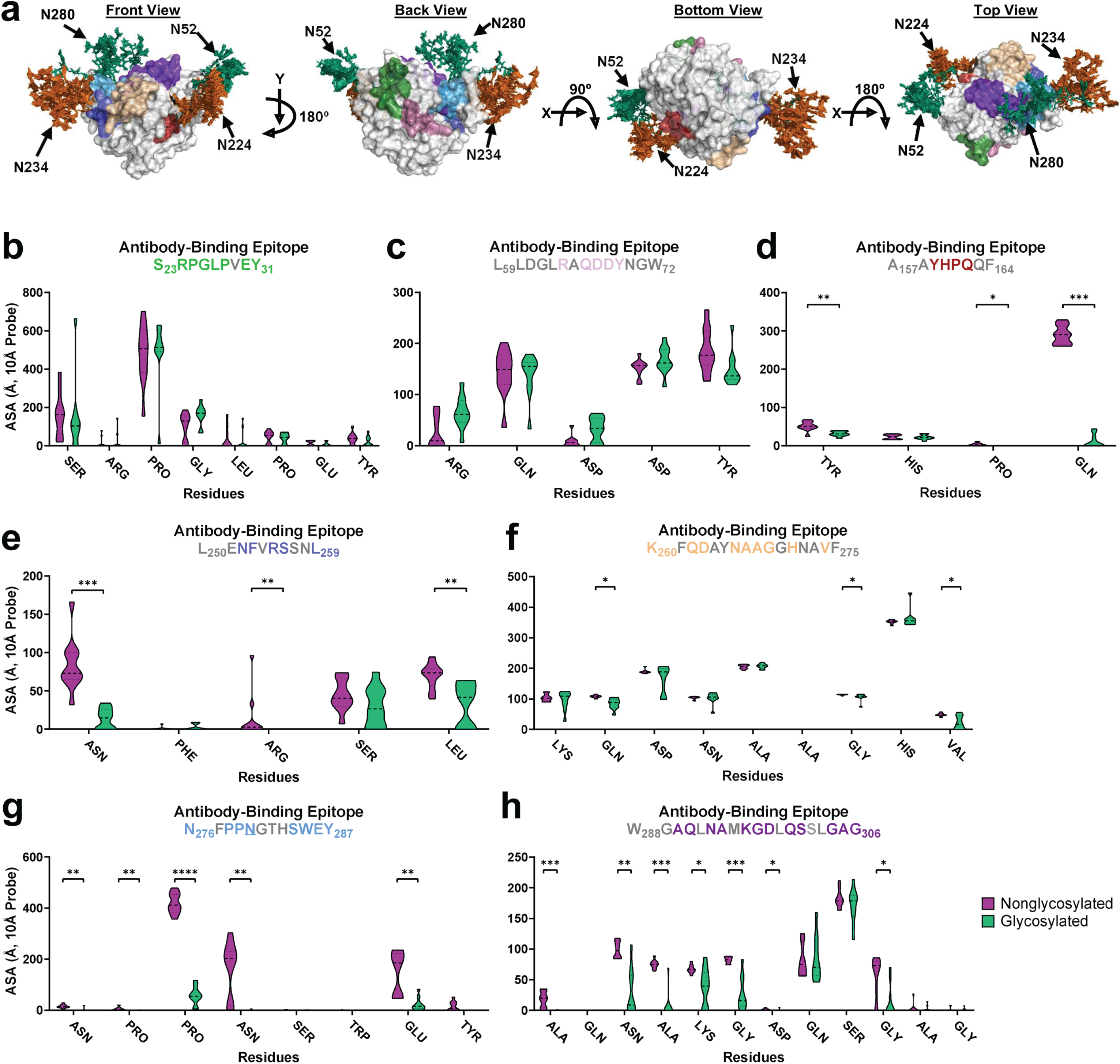
Molecular dynamics simulations illustrate the impact of Ag85B N-glycosylation on the accessibility of known antibody binding epitopes. **(a)** Hierarchical ensemble representation of the glycomics-informed glycopeptidomics site-specific abundant glycan structures of Ag85B generated after three 1 μs-long molecular dynamics simulations viewed from front, back, bottom, and top (from left to right), displaying specific antibody binding epitope locations distinguished through color coding. Antibody accessible surface areas of various antibody epitopes identified by Rinaldi et al. 2018 using proteolytic affinity-mass spectrometry and biosensor analysis displayed through violin plots for **(b)** site S23-Y31 (green on panel a), **(c)** site L59-W72 (pink on panel a), **(d)** site A157-F164 (red in panel a), **(e)** site L250-L259 (blue in panel a), **(f)** site K260-F275 (tan in panel a), **(g)** site N276-Y287 (light-blue in panel a), and **(h)** site W288-G306 (purple in panel a). Identified N-glycosylated asparagines are underlined in the graph title. For clarity, amino acids where there was no accessibility before and after adding glycans at identified N-glycan sequons are colored in gray in the graph titles and are removed from the shown data on the x-axis. ∗∗∗∗*p* < 0.0001, ∗∗∗*p* < 0.001, ∗∗*p* < 0.01, and ∗*p* < 0.05 represent significant differences between groups determined using either Welch’s t-test or the Mann-Whitney U test, depending on distribution normality.

### Molecular dynamics simulations illustrate the detrimental effect of Ag85B N-glycosylation on established antibody binding sites

Humoral immunity is a key mediator by which vaccines achieve “sterilizing" immunity against pathogens (44). One common immune evasion strategy employed by pathogens is the use of “glycan shields,” where glycans sterically obstruct antibody binding by masking surface-exposed epitopes (45, 46). Hence, to investigate whether glycosylation similarly impairs antibody access, we utilized previously identified antibody binding sites against Ag85B (47). These sites were identified using proteolytic affinity-mass spectrometry and biosensor analysis against human serum samples from healthy, vaccinated, and infected donors.

The identified antibody-binding sites were mapped onto the glycosylated Ag85B ensemble structure generated from the molecular dynamics simulations **(Fig. 3a)**. When the antibody binding sites were located distantly from glycosylation sites, little to no loss in accessibility to a 10 Å probe was observed, as demonstrated by sites S23-Y31 and L59-W72 **(Fig. 3b-3c)**. However, as antibody sites approached glycosylated residues, greater loss of accessibility was observed, with several residues within each site exhibiting a significant reduction in accessibility to the 10 Å probe **(Fig. 3d-3h)**.

Notably, antibody binding site K260-F275, although not directly adjacent to a glycosylation site, contains at least three residues with a significant loss of accessibility **(Fig. 3f)**. In addition, glycosylation site N280 is located within antibody binding site N276-Y287, where many residues show a pronounced decrease in surface accessibility. Compared to sites A157-F164 (21.6–97.3% loss of accessibility) and K260-F275 (4.3-63.6% loss), both of which are not directly at a glycosylation site, site N276-Y287 exhibits a far more extensive loss, ranging from 81.3 to 99.9% **(Fig. 3g)**. Antibody binding site W288–G306 is also significantly affected by the nearby N280 glycan even though it does not contain a glycosylation site within the epitope. Most exposed surface residues in this site lose accessibility to a 10 Å probe, with reductions ranging from 45.8 to 99.8% **(Fig. 3h)**. The significant loss of accessibility observed across multiple antibody binding sites, either in proximity to or directly overlapping glycosylation sites, highlights the substantial role glycans play in shielding key epitopes from recognition by the adaptive immune response.

### Glycosylation of Ag85B negatively impacts antigenicity

Using *in silico* approaches, we predict that glycosylation significantly impairs the accessibility of the antibody-binding sites on Ag85B. We evaluated the effect *in vitro* by using a monoclonal antibody targeting the Ag85 complex to assess whether glycosylation impacts antibody binding. We performed a competitive ELISA in which plates were coated with bacterially expressed Ag85B, and the ability of Ag85B variants to compete for antibody binding was evaluated. We found that glycosylation of Ag85B significantly reduces its ability to inhibit antibody binding to the bacterial protein **(Fig. 4a)**.

**Figure 4:**
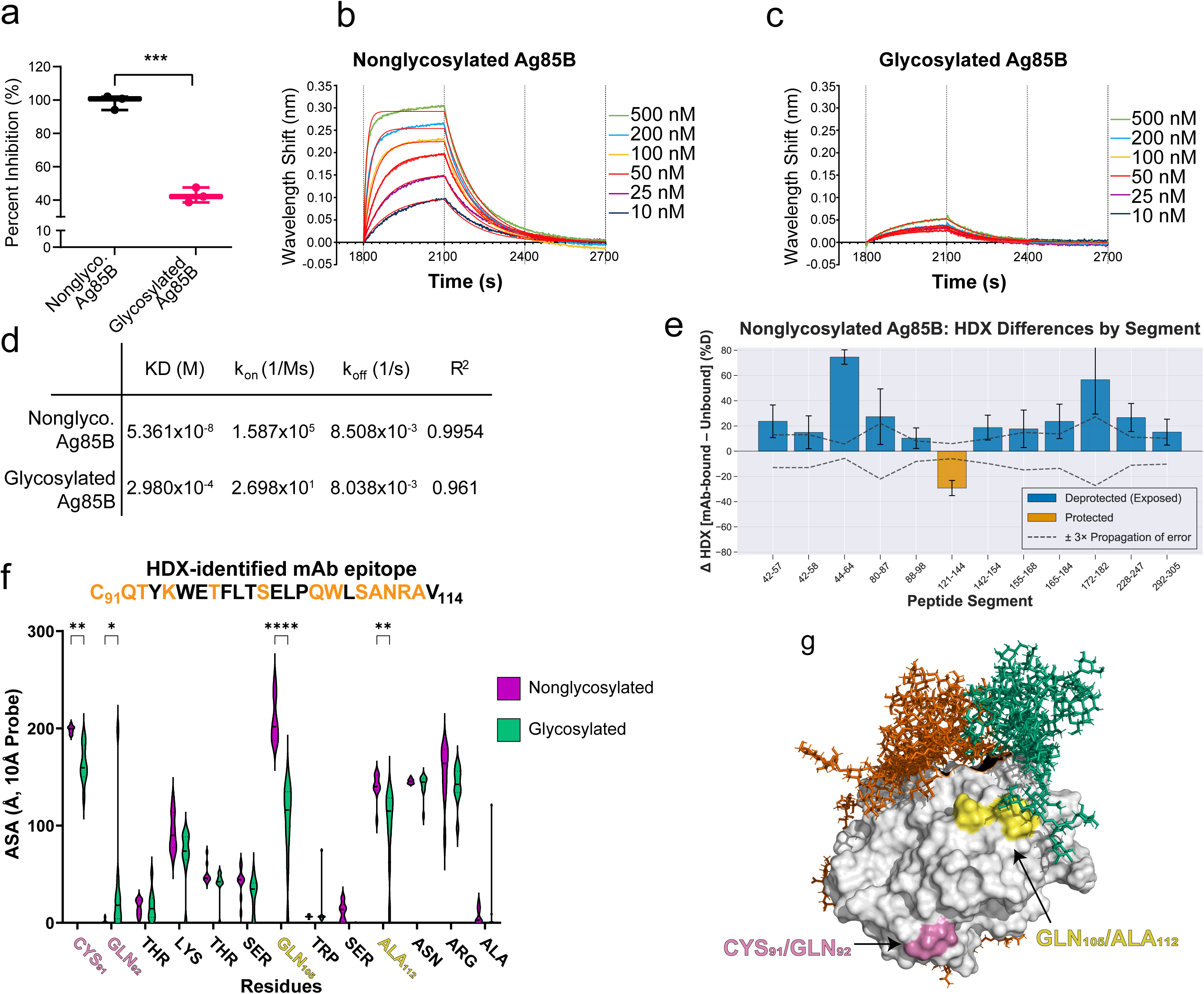
Glycosylation of Ag85B impacts its antibody binding. **(a)** Glycosylation of Ag85B impairs binding of a monoclonal antibody against the Ag85 Complex (BEI Resources, Clone P4B3G2) compared to non-glycosylated Ag85B, as determined by competitive ELISA. Values represent the mean ± SD of the percentage of inhibition. ∗∗∗∗p < 0.0001, ∗∗p < 0.01, and ns represent significant differences between groups determined using two-way ANOVA with Tukey’s multiple comparisons test. Biolayer interferometry using AMC anti-mouse Fc Biosensors loaded with a monoclonal antibody against Ag85 Complex (from panel a) binding association and dissociation to **(b)** nonglycosylated Ag85B and **(c)** glycosylated Ag85B. **(d)** KD, kon, koff, and R2 were determined by global 1:1 fitting analysis using the Sartorius Octet Analysis Suite. **(e)** Epitope mapping of mAb P4B3G2 against non-glycosylated Ag85B using differences in HDX rates, data averaged from four timepoints. A positive value indicates greater accessibility in the bound state, while a negative value indicates reduced accessibility in the bound state. Peptides with exchange differences <3x propagation of error omitted for clarity. **(f)** Antibody accessible surface areas of residues within the HDX-identified epitope. ∗∗∗∗p < 0.0001, ∗∗∗p < 0.001, ∗∗p < 0.01, and ∗p < 0.05 represent significant differences between groups determined using either Welch’s t-test or the Mann-Whitney U test, depending on distribution normality. **(g)** 3D representation of HDX-identified epitope with residues of interest highlighted.

To further characterize these interactions, we employed biolayer interferometry (BLI) to measure the binding kinetics of the monoclonal antibody against the different Ag85B variants **(Fig. 4b-4e)**. The bacterially expressed Ag85B, representing the wild type binding profile expected to be seen in natural infections, exhibited a robust wavelength shift of approximately 0.30 nm during association, which returned to baseline upon dissociation **(Fig. 4b)**. In contrast, the glycosylated Ag85B showed a substantially reduced shift of about 0.05 nm, indicating a decreased association rate (**Fig. 4c**). Bacterial Ag85B had a dissociation constant of 54 nM, while glycosylated Ag85B had a dramatic reduction in binding affinity, with a dissociation constant of 298 μM (**Fig. 4d**).

To determine whether the reduced binding affinity results from glycosylation limiting antibody access, we used hydrogen-deuterium exchange mass spectrometry (HDX-MS) to map the epitope of our mAb to a specific region of the protein. HDX-MS identified a single region that became protected from deuterium exchange upon mAb binding, indicating that this site is the most likely antibody epitope **(Fig. 4e)**. Glycosylated Ag85B was omitted from analysis due to poor binding. Consistent with our hypothesis, the solvent-accessible surface areas (ASAs) of several residues within this epitope, particularly Gln_105_ and Ala_112_, were significantly reduced in the glycosylated model **(Fig. 4f)**. Interestingly, Gln_92_ showed increased ASA in the glycosylated model; together with reduced accessibility of the neighboring Cys_91_ residue, this suggests that glycosylation may induce a conformational change that exposes residues that are otherwise shielded. Supporting this, when represented as a 3D model **(Fig. 4g)**, it becomes clear that Cys_91_ and Gln_92_ are distal to the glycans, whereas Gln_105_ and Ala_112_ are proximal.

### N-glycans are positioned to hinder the processing/presentation of known T cell epitopes of Ag85B

Similar to how unnatural mammalian host glycans can shield antibody binding regions on the native protein, they can also impact cellular adaptive immunity through several mechanisms: (1) by masking protease cleavage sites and preventing proper processing of MHC epitopes, (2) by interfering with or altering MHC molecule binding in the microsomes, (3) through the presentation of glycosylated peptides to T cells, which can modulate T cell responses differently from the native peptides, and (4) by inducing deamidation of asparagine to aspartic acid during antigen processing, leading to the presentation of epitopes that differ from those derived from the native protein (48, 49).

To assess how glycosylation affects cellular immunity, we mapped previously identified MHC-binding epitopes onto the glycosylated Ag85B ensemble structure generated from molecular dynamics simulations **(Fig. 5a)** (50). We hypothesized that sites proximal to non-native N-glycosylation would exhibit reduced accessibility. For example, the peptide A111–E125, identified as presented by DPB1*04:01 during human latent TB infection (51), is not located near any glycosylation sites **(Fig. 5a)**. When examining its accessible surface area using a 10 Å probe, most residues in this peptide exhibit minimal variation **(Fig. 5b)**, with two residues showing significant but mild changes in accessibility (C113 decreasing, A114 increasing). In contrast, F121–P138, another T cell peptide identified during latent TB infection (52, 53), lies near two glycosylation sites (N52 and N224). Several of its surface-accessible residues exhibit a substantial decrease in accessibility upon glycosylation, ranging from 20.6 to 99.5% **(Fig. 5c)**. Unlike A111–E125, no residues within F121–P138 display increased accessibility. These data suggest that proximity to non-native N-glycosylation may hinder epitope accessibility.

**Figure 5:**
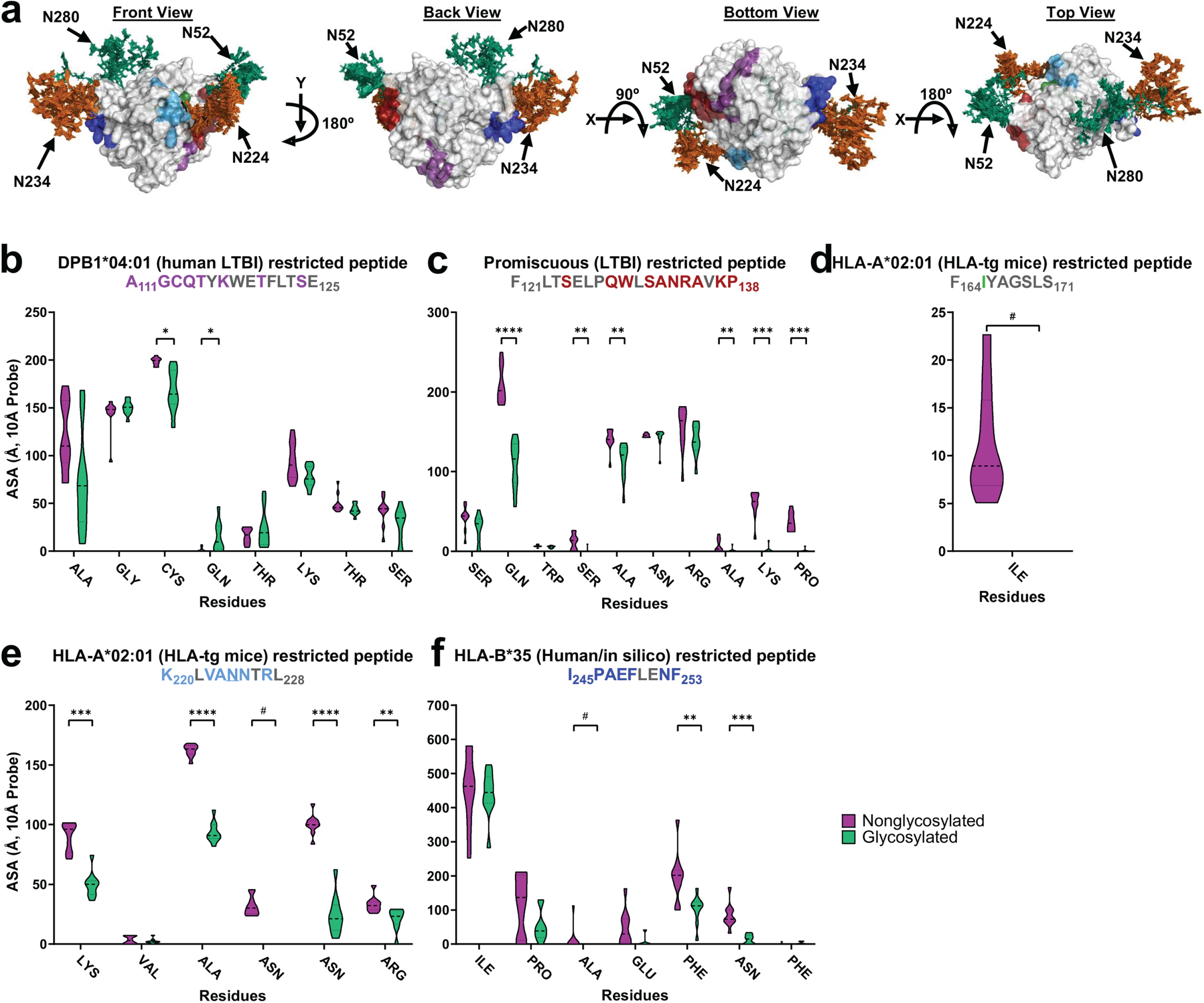
Molecular dynamics simulations demonstrate the impact of Ag85B N-glycosylation on the accessibility of known T-cell epitopes. **(a)** Hierarchical ensemble representation of the glycomics-informed glycopeptidomics site-specific abundant glycan structures of Ag85B generated after three 1 μs-long molecular dynamics simulations viewed from front, back, bottom, and top (from left to right), displaying specific T-cell epitope locations distinguished through color coding. Accessible surface areas of various T-cell epitopes displayed through violin plots for **(b)** site A111-E125 (purple on panel a), **(c)** site F121-P138 (red on panel a), **(d)** site F164-S171 (green on panel a), **(e)** site K220-L228 (light-blue on panel a), and **(f)** site I245-F252 (blue on panel a). Identified N-glycosylated asparagines are underlined in the graph title. For clarity, amino acids that were previously inaccessible and remained so after adding glycans at identified N-glycan sequons are colored gray in the graph titles and removed from the shown data on the x-axis. ∗∗∗∗*p* < 0.0001, ∗∗∗*p* < 0.001, ∗∗*p* < 0.01, and ∗*p* < 0.05 represent significant differences between groups determined using either Welch’s t-test or the Mann-Whitney U test, depending on distribution normality. # represents complete loss of accessibility after glycan addition.

Two additional Ag85B peptides, F164–S171 and K220–L228, were identified using human MHC transgenic mice and are presented to T cells via HLA-A*02:01 in a DNA vaccination model **(Fig. 5d–5e)** (54). Peptides presented by HLA-A*02:01 are of particular interest due to the high prevalence of this haplotype in the global population (55, 56). Glycosylation of Ag85B significantly affects the accessibility of both peptides. For F164–S171, only one residue is surface-accessible, and this accessibility is completely lost upon the addition of a glycan at N224 **(Fig. 5d)**. In comparison, K220–L228 contains multiple accessible residues, many of which show a loss of accessibility ranging from 41.4 to 100% when N224 is glycosylated **(Fig. 5e)**.

Finally, using a reverse immunogenetics approach, the peptide I245–F253 was identified as an HLA-B*35-restricted epitope in humans (57). Within this peptide, several residues, particularly those near the N234 glycan, exhibit a marked reduction in surface accessibility, ranging from 45.6-100% **(Fig. 5f)**. Together, the substantial loss of accessibility observed in several T cell epitopes, whether adjacent to or directly overlapping with glycosylation sites, demonstrates the potential for glycans to interfere with epitope processing and presentation. In some cases, glycopeptides may be presented in place of the native epitope, with important implications for T cell recognition and immunogenicity.

### Glycosylation of Ag85B inhibits cellular immune response

Based on our predictions that proximal glycans may shield T cell epitopes (**Fig. 5**), we conducted an *ex vivo* T cell proliferation assay (**Fig. 6a**) to examine how glycosylation influences T cell stimulation by Ag85B. Human peripheral blood mononuclear cells (PBMCs) were isolated from BCG-vaccinated healthy donors and then stimulated with equal amounts of either glycosylated or nonglycosylated Ag85B. An increase in CD4^+^ T cell proliferation was observed in both groups compared with media alone, confirming that both protein forms retain immunostimulatory capacity. However, glycosylated Ag85B induced a significantly reduced proliferative response relative to the nonglycosylated counterpart (**Fig. 6b**), indicating that glycosylation dampens antigen-specific CD4+ T cell activation. These findings are consistent with our structural predictions and suggest that surface glycans may partially mask immunodominant epitopes or otherwise interfere with effective antigen processing and/or presentation.

**Figure 6:**
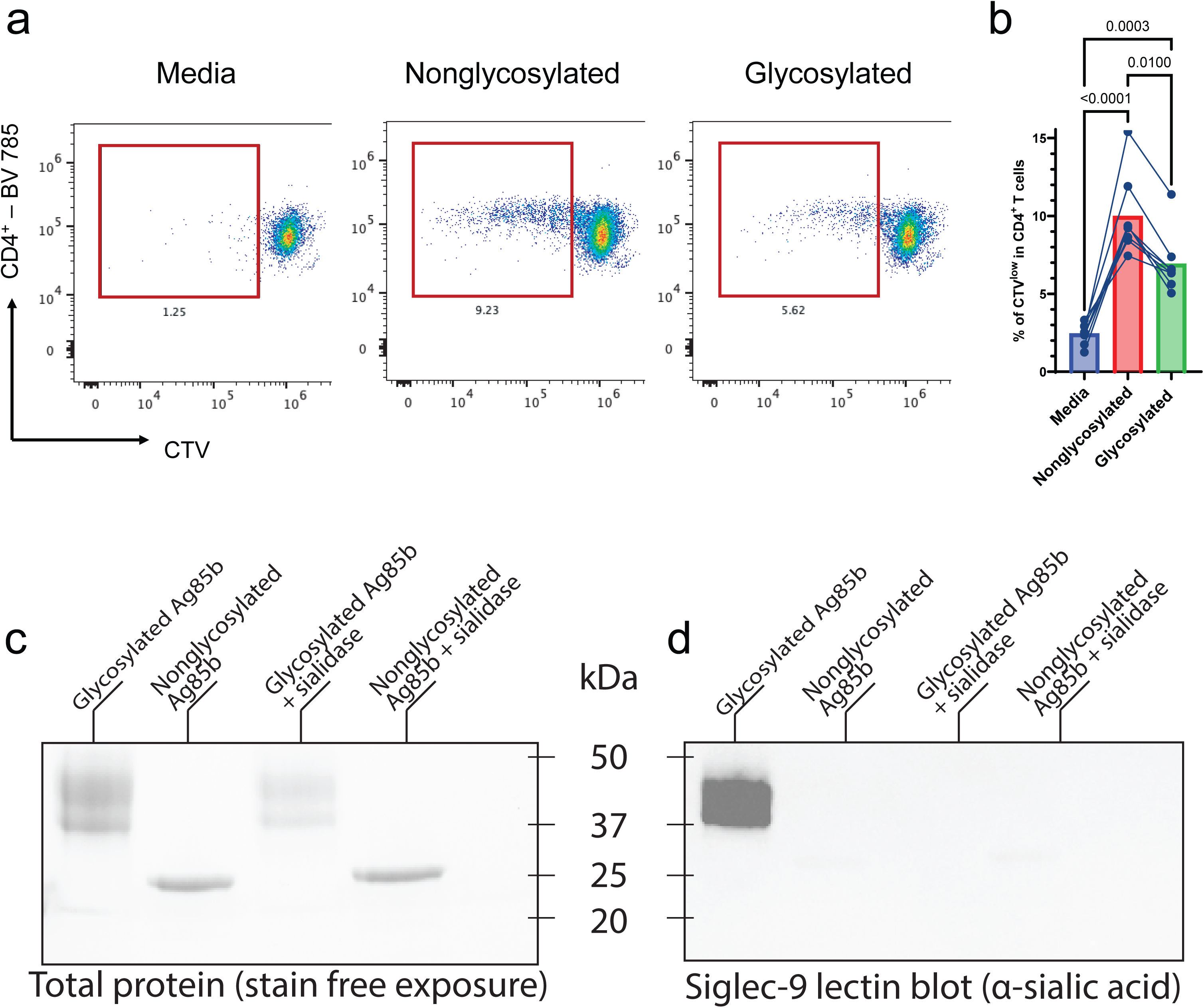
Glycosylation of Ag85B impedes T cell proliferation and promotes immunoinhibitory Siglec binding. **(a)** Human PBMCs were stimulated with Ag85B variants, and helper T cell proliferation was assessed by CellTrace™ Violet dye dilution by flow cytometry. Helper T cells were assessed using CD4 gating, and proliferation was identified by gating on CTV_low_ populations. (**b**) Quantitation of CTVlow CD4+ T cell populations among stimulation groups. Significant differences between groups were determined using two-way ANOVA with Tukey’s multiple comparisons test. Stain-free gel image (**c**) and Siglec-9 lectin blot analysis (**d**) of nonglycosylated and glycosylated Ag85B with or without sialidase treatment.

Building on our glycomics and glycoproteomics analyses, which revealed extensive sialylation of glycans on mammalian-expressed Ag85B, we next examined whether this modification could mediate interactions with immunoinhibitory lectins. As a proof of concept, we focused on Siglec-9, an inhibitory siglec known to recognize a range of sialoglycan structures, with a reported preference for sialyl Lewis X motifs (58, 59). Siglec-9 plays a key role in inhibiting immune responses in a variety of diseases, including viral infection (60) and cancer (61). We assessed Siglec-9 binding capability with a lectin blot **(Fig. 6c-d)**. The Expi293-expressed (glycosylated) Ag85B bound to Siglec-9, but the BL21-expressed Ag85B (non-glycosylated) did not, demonstrating that Siglec-9 recognition is glycan-dependent. Binding to glycosylated Ag85B was abrogated upon treatment with a broadly-acting α-2,3/6/8/9 sialidase **(Fig. 6c-d)**, confirming that terminal sialic acids mediate this interaction.

Together, these data support a model in which mammalian glycosylation of Ag85B not only reduces CD4+ T cell proliferation but also introduces sialylated ligands capable of engaging inhibitory siglec receptors. This dual mechanism, epitope masking and active engagement of inhibitory lectins, may contribute to the attenuated cellular immune response observed with glycosylated Ag85B.

## Discussion

The success of COVID-19 mRNA vaccines has highlighted the promise of nucleic acid vaccine technologies as effective preventive tools. This breakthrough has generated considerable interest in extending mRNA and DNA vaccine platforms to a wide range of infectious diseases. Nonetheless, although nucleic acid vaccines can be well-suited for viral antigens produced naturally within host cells, their use against non-viral pathogens like bacteria and parasites involves distinct biological challenges. Notably, when these foreign antigens are expressed in mammalian cells, they may acquire host-derived post-translational modifications (PTMs), particularly N-linked glycosylation, that are absent in their native pathogen context. These glycan modifications can mask critical epitopes or engage immune regulatory pathways, ultimately dampening the desired immune response. Such unintended alterations could undermine the efficacy of nucleic acid vaccines targeting non-viral pathogens.

Importantly, the role of glycosylation in shaping immune responses is not limited to non-viral pathogens (62, 63). In viral vaccine targets such as the HIV-1 envelope glycoprotein (Env), dense glycan shielding on the gp120 subunit can hinder antibody access to conserved neutralizing epitopes (64, 65). Selective removal of N-glycosylation sites unmasks these epitopes, thereby directing the immune system to generate broadly neutralizing antibodies (43, 66, 67). Similarly, in SARS-CoV-2, the spike protein’s glycosylation landscape influences antigenicity and immune targeting (68, 69). Eliminating specific glycosylation sites on the spike can enhance the breadth and potency of the antibody response (70–72).

Findings across both viral and non-viral systems highlight that glycosylation is not a peripheral concern but a central determinant of vaccine efficacy. As such, nucleic acid vaccine design must move beyond sequence selection to consider how expression in mammalian hosts alters antigen structure and immune presentation. Incorporating glycosylation-aware design strategies will be critical to unlocking the full potential of nucleic acid vaccines across diverse pathogen classes.

Our investigation of the *Mtb* antigen Ag85B provides a direct, mechanistically informative example of this principle. Ag85B is a secreted, highly immunogenic protein critical for *Mtb* cell wall biosynthesis and has been widely used in TB vaccine candidates, including those delivered via nucleic acid platforms (19, 21–23, 26). In its native bacterial context, Ag85B is not glycosylated. However, when expressed in mammalian host cells during nucleic acid vaccination, Ag85B acquires N-linked glycans at specific asparagine residues. We demonstrated that this aberrant glycosylation significantly reduces its antigenicity and dampens immune activation.

At a structural level, these glycans likely occlude key B-cell or T-cell epitopes, hinder antigen processing by antigen-presenting cells, or alter MHC loading and presentation, mechanisms well-documented in other glycan-shielded antigens such as HIV Env (62, 64, 67, 73). Glycans can also engage inhibitory immune receptors (e.g., DC-SIGN, siglecs), actively suppressing immune responses (32, 74, 75). Understanding these molecular mechanisms, including how host-derived glycosylation alters epitope exposure, antigen stability, and receptor interactions, is critical not only for optimizing individual antigens but for establishing design principles that guide antigen engineering across vaccine platforms. A previous study demonstrated the power of informed protein engineering in glycoconjugate vaccine design by utilizing glycan conjugation on Ag85B via lysine residues. Targeted mutations prevented conjugation at key epitopes, preserving T cell function while maintaining glycan conjugation at non-critical sites (76).

While N-linked glycosylation is a well-characterized and structurally defined post-translational modification with clear implications for antigen recognition and immune activation, O-linked glycosylation may also modulate immune responses. O-glycans, typically added to serine or threonine residues, can influence protein conformation, mucosal immunity, and receptor engagement (32, 77). However, O-linked glycosylation is inherently more heterogeneous and less predictable than N-linked glycosylation, with no strict consensus sequence and extensive variation in glycan composition and site occupancy (77). This lack of predictability makes both identifying relevant O-glycosylation sites and modulating them technically challenging. Therefore, prioritizing O-glycosylation for immunogen engineering may not be practical unless N-glycan modulation fails to restore immunogenicity. However, characterization of potential O-glycosylation sites via approaches such as mass spectrometry or lectin blots may be an important yet overlooked step in the characterization of engineered antigens. In this work, we specifically focus on N-linked glycosylation due to its defined structural rules, clearer immunological impact, and greater potential for rational design in nucleic acid vaccine contexts.

In this study, we provide mechanistic insight into how host-cell-mediated N-glycosylation impairs the antigenicity of Ag85B, a major TB vaccine candidate antigen. When expressed in human Expi293 cells, Ag85B is microheterogeneously glycosylated at four canonical N-linked glycosylation sequons, leading to altered structural and immunological properties. We demonstrated, using molecular dynamics simulations, that these glycans significantly impede the surface accessibility of Ag85B, including the occlusion of known B-cell and T-cell epitopes, thereby hindering immune recognition. Functional assays further confirmed that glycosylated Ag85B shows reduced binding to an Ag85B-specific monoclonal antibody, as measured by ELISA, BLI, and HDX MS. Additionally, glycosylation affects cellular immunity by reducing T cell activation and binding to the immunoinhibitory Siglec-9. Therefore, understanding how the host influences antigen remodeling highlights the importance of redesigning nucleic acid vaccines. These vaccines should comprehensively incorporate biochemical and immunological factors to prevent immune silencing caused by glycans.

This study has broad implications for nucleic acid vaccine design, particularly for non-viral pathogens. As nucleic acid platforms drive antigen expression in mammalian cells, unintended post-translational modifications, particularly N-linked glycosylation, may structurally remodel microbial proteins, thereby diminishing their immunogenic potential. Our work emphasizes the importance of evaluating and, when necessary, modifying candidate antigens to preserve epitope integrity and immune visibility. Overall, our results emphasize the importance of including glycosylation-aware design principles in future nucleic acid vaccine development, highlighting the role of structural and molecular immunology in connecting antigen expression to immune activation.

## Methods

### Proteins and Bacteria

*Mycobacterium tuberculosis* H37Rv strain was acquired from the Biodefense and Emerging Infections (BEI) Research Resources Repository of the National Institute of Allergy and Infectious Diseases at the National Institutes of Health (NIH). Native Ag85B proteins were purified from the *Mtb* strain H37Rv.

### Expression and purification of *E. coli* BL21-expressed recombinant Ag85B

Plasmids with the Ag85B gene were generated using polymerase chain reactions (PCR) and DNA assembly techniques (NEBuilder HiFi DNA Assembly, New England Biolabs) to insert the gene into a pET-28b(+) vector backbone with an N-terminal His-tag and TEV cleavage site. All PCR reactions were confirmed by gel electrophoresis using a 1% agarose gel in Tris-acetate-EDTA buffer. PCR reactions were purified using the Invitrogen PureLink Quick PCR Purification Kit.

Plasmids were transformed into *Escherichia coli* (*E. coli*) BL21 cells and incubated at 37°C with an orbital shaking speed of 200 rpm until an OD of 0.7-1.0. Protein expression was induced with 1 mM isopropyl β-D-thiogalactopyranoside (IPTG) and incubated at 18°C with an orbital shaking speed of 200 rpm for 24 hours before harvesting cells for protein purification. All proteins were purified using IMAC columns (BioRad, Cat. # 12009287) and then separated by a HiLoad Superdex 200 pg preparative size-exclusion column (Cytiva, Cat. # 28989335) using a BioRad fast-protein liquid chromatography. Proteins were validated using mass spectrometry.

### Expression and purification of mammalian-expressed recombinant Ag85B

Plasmids with the Ag85B variant gene insertions were first generated for mammalian cell protein expression. *Homo sapiens* codon-optimized genes with the tissue plasminogen activator leader signal sequence at the N-terminal and a TEV cleavage site with an 8xHis tag at the C-terminal were synthesized by Twist Biosciences. 293-F Ag85B synthesized gene was inserted into a pGEc2-DEST vector with a CMV promoter using PCR and DNA assembly technique (NEBuilder HiFi DNA Assembly, New England Biolabs). All PCR reactions were confirmed by gel electrophoresis using a 1% agarose gel in Tris-acetate-EDTA buffer (Addgene). PCR reactions were purified using the PCR purification methods (Invitrogen PureLink Quick PCR Purification Kit). Mutant variant genes were cloned through Twist Biosciences. All plasmids were confirmed through sequencing methods (Eurofins).

Recombinant Ag85B proteins were expressed in Expi293F™ cells cultured in Expi293 expression medium (Gibco, Cat. # A14635). Cells were transiently transfected with the ExpiFectamine™ 293 Transfection Kit (Gibco, Cat. # A14524) or Polyethylenimine “Max” (PEI MAX) (Polysciences, CAS # 49553-93-7) following the manufacturer’s protocol or an established protocol by Dr. Dumoux, respectively. All proteins were purified using IMAC columns (BioRad, Cat. # 12009287) and then separated by a HiLoad Superdex 200 pg preparative size-exclusion column (Cytiva, Cat. # 28989335) using a BioRad fast protein liquid chromatography. Proteins were validated using mass spectrometry.

### Released Ag85B N-glycan analysis (glycomics) via LC-MS

Glycans from 20µg purified recombinant Ag85B were released and labeled with InstantPC dye using the Agilent AdvanceBio Gly-X N-glycan prep with InstantPC kit (Agilent, Part #: GX96-201PC) as described in the kit instructions. 2µL of the resulting 100µL sample was injected onto a 2.1×150mm 1.8µm-particle size AdvanceBio Amide HILIC column (Cat. #: 859750-913). Glycans were separated with a two-buffer LC gradient using 50 mM ammonium formate as buffer A and acetonitrile with 0.1% formic acid as buffer B with an Agilent 1290 Infinity II liquid chromatography system. The buffer gradient proceeded as follows: 0-45min 23 to 44% A, 45-46min 44 to 56% A, 46-47min 56 to 60% A, 47-49min 60 to 23% A, 49-60min 23% A. Eluting glycans were analyzed with an Agilent 6545XT quadrupole time-of-flight (QTOF) mass spectrometer with a dual Agilent Jet Stream (AJS) electrospray ionization (ESI) source operating in positive ion mode. Raw files were analyzed utilizing Skyline with a transition list derived from the human N-glycan compositions described in the GlyGen(78) Database (accessed August 2024). Glycan compositions were relatively quantified by summing the peak area of all observed adducts and isomers using Skyline(79) and Python. Figures were generated using GraphPad Prism 9 software.

### N-linked glycopeptide and site occupancy analysis – Sample Preparation

Intact glycopeptides were generated utilizing purified Ag85B. 50 µg of purified glycoprotein was reduced for 1 hour at 56°C in 10 mM DTT. Ag85B cysteines were alkylated using 25 mM iodoacetamide in the dark at room temperature for 45 minutes. Alkylation was quenched with an additional 10 mM DTT. Glycoprotein was then digested with one of two separate protease conditions: 1) α-lytic protease (NEB, Part #: P8113S), or 2) trypsin/LysC mix (Promega, Part #: V5071) followed by GluC (Promega, Part #: V1651). All protease digestions were carried out at 37°C overnight (∼16-18 hours) at a 1:20 µg:µg protease:protein ratio (e.g., protease condition 2 required two overnight digests). After each digest, the protease was heat-inactivated at 95°C for 5 minutes.

Glycopeptides were desalted via reverse-phase cleanup using a BioPureSPN MINI C18 column (The Nest Group, Part #: HUM S18V). Desalted peptides were dried down with a vacuum centrifuge and then resuspended in a solution of 95.9% water, 4% acetonitrile, and 0.1% formic acid. Resuspended peptides were filtered through a conditioned 0.2µm filter (PALL, Cat. #: ODM02C34) before transferring to autosampler vials. For site occupancy analysis(29), desalted glycopeptides were instead resuspended in 50µL of a 50 mM sodium phosphate buffer with 5µL/500 units EndoH (NEB, Part #: P0702S) at 37°C overnight, followed by the addition of 50µL 1% trifluoroacetic acid (TFA) to stop the reaction. Partially deglycosylated peptides were then subjected to another round of reverse phase cleanup (see above), then resuspended in 50µL of a 20 mM ammonium bicarbonate in heavy-labeled ^18^O water (Cambridge Isotope Laboratories, Part #: DLM-4-100). Glycopeptides were treated with 2µL/20 units PNGaseF (Promega, Part #: V4831) overnight at 37°C. Deglycosylated peptides were then subjected to a final round of reverse-phase cleanup and 0.2µm filtering (see above) before transfer to autosampler vials.

### N-linked glycopeptide and site occupancy analysis – LC-MS/MS Analysis

For intact glycopeptide analysis, 1µg of Ag85B glycopeptides was injected onto a 75µmx15cm Acclaim PepMap 100 C18 nanoViper column (Thermo Fisher Scientific, Part #: 164940) and eluted into the nanospray source of an Orbitrap Eclipse Tribrid mass spectrometer with a field asymmetric ion mobility spectrometry (FAIMS) frontend interface (Thermo Fisher Scientific). Glycopeptides were separated with a two-buffer LC gradient using water with 0.1% formic acid as buffer A and 80% acetonitrile with 0.1% formic acid as buffer B utilizing a Vanquish liquid chromatography system (Thermo Fisher Scientific) with a flow rate of 0.4 µL/min. The 90-minute buffer gradient proceeded: 0-6min 2.5% B, 6-76min 2.5 to 32.5% B, 76-88min 32.5% to 100% B, followed by a 1 min column wash with 100% B at 1 µL/min. The spray voltage was set to 2.35 kV, and the ion transfer tube was heated to 305°C. FAIMS compensation voltages were set to cycle through −35 V, −40 V, and −50 V. Full MS scans were acquired from m/z 200 to 2000 at 60,000 resolution in the Orbitrap. MS/MS scans were triggered with an intensity threshold of 2 × 10^4^ and a mass tolerance of 20 ppm. MS/MS scans were acquired at a resolution of 15,000 in the Orbitrap via stepped higher collisional dissociation (stepHCD), with HCD collision energies of 15%, 25%, and 35% and an isolation window of 1.6 m/z.

For site occupancy analysis (29, 63), 10µg of EndoH/PNGaseF-treated peptides were injected onto a 2.1×150mm 2.7µm pore size Peptide Mapping C18 column (AdvanceBio, Part #: 653750-902) eluted into the dual Agilent Jet Stream (AJS) electrospray source of an Agilent 6560 mass spectrometer (Agilent). Peptides were separated with a two-buffer LC gradient using water with 0.1% formic acid as buffer A and acetonitrile with 0.1% formic acid as buffer B utilizing a 1290 Infinity II liquid chromatography system (Agilent) with a flow rate of 0.4 mL/min. The 45-minute buffer gradient proceeded as follows: 0-2.5min 2.5% B, 2.5-4min 2.5 to 6% B, 4-12min 6% to 22% B, 12-33min 22 to 29% B, 33-38min 29 to 34% B, 38-40min 34 to 81% B, 40-43min constant 81% B wash, then equilibration from 43-45min 81 to 2.5% B. Eluting glycans were analyzed with an Agilent 6560 (QTOF) mass spectrometer equipped with a dual AJS ESI source operating in QTOF-only positive ion mode. MS1 full scans were acquired from 100-3000 m/z, with an acquisition rate of 8 spectra/second and 125ms/spectrum. Data-dependent MS/MS scans were triggered with an isolation window of 1.3 m/z at a rate of 3 spectra/second and 333 ms/spectrum.

### N-linked glycopeptide and site occupancy analysis – Data Analysis

For intact glycopeptide analysis, the resulting RAW files were searched for glycopeptide spectral matches (GPSMs) using the search engine pGlyco3 (80). RAW files were searched against a FASTA database containing Ag85B and common contaminants and a custom glycan database derived from released glycan compositions identified in glycomics experiments (see above). Search was conducted with a precursor tolerance of 10 ppm, fragment tolerance of 20 ppm, and glycopeptide FDR set to 0.01. Resulting glycopeptide spectral matches (GPSMs) were manually inspected to ensure accuracy. GPSMs were processed by the ppmFixer script (81) to adjust for common errors due to monoisotopic peak misidentification, and site-specific glycan compositions were relatively quantified using spectral counts (82).

Agilent .d file folders were analyzed for site occupancy analysis using MassHunter Qualitative Analysis (Agilent). Extracted ion chromatograms for predicted doubly and triply charged tryptic and α-lytic peptides were generated, and peak areas were obtained. The unmodified peptide denotes the non-glycosylated fraction, the +203 modification denotes high-mannose and hybrid structures, and the +3 modification denotes complex structures. Peak areas for all observable predicted peptides and charge states were summed to obtain the final site-specific glycoprofile. Figures were generated using GraphPad Prism 9 software.

### 3D Structural Modeling and Molecular Dynamics Simulations of Nonglycosylated and Glycosylated Ag85B

#### Mtb Ag85B protein structure and glycoform generation

An experimentally determined X-ray diffraction structure of *Mtb* Ag85B (RCSB PDB: 1F0N) was obtained through the RCSB Protein Data Bank (40, 83). Glycans were selected to install glycosylated Ag85B N-glycan sequons based on glycomics-informed glycoproteomics analysis of site-specific abundant glycoforms. The glycoprotein builder available at GLYCAM-Web (www.glycam.org) was employed together with an in-house method that adjusts asparagine side chain torsion angles and glycosidic linkages to relieve atomic overlaps with the core protein backbone, as described previously (36, 38).

#### Energy minimization and Molecular Dynamics (MD) Simulations

Each protein structure was placed in a periodic box of OPCBOX water molecules with a 1.4 Å buffer between the solute and solvent box. Additionally, 66 sodium and 60 chlorine ions were placed randomly within the box to mimic physiological salt concentrations (∼150 mM NaCl). Energy minimization for each production was completed following a previously established ten-step protocol involving 4000 total steps of minimization and 40,000 steps of MD, which constitute a series of energy minimizations and relaxation steps that allow the system to relax gradually under a constant pressure of 1 atm and a constant temperature of 300 K (84). All MD simulations were performed in the NVT canonical ensemble with a CUDA-enabled implementation of PMEMD included in the Amber24 software suite (85–88). The ff19SB force field and GLYCAM06j force field were used for the protein and carbohydrate moieties, respectively. The Bussi thermostat was used to maintain a constant temperature of 300 K. Periodic boundary conditions under constant-volume, no-pressure regulation, and a 10 Å cutoff for non-bonded interactions were employed throughout the simulations. The SHAKE algorithm was used to constrain bonds with hydrogen atoms, and the integration time step was set to 2 fs. Each system was then equilibrated for 10 ns before initiating 3 independent 1-µs production MD simulations with random starting velocities, for a total of 3 µs per system.

#### Surface Area Accessibility Analysis

To determine solvent-accessible surface areas, 3D structures of each frame (every 1 ns) from each production run were subsequently analyzed for their ability to interact with a 1.4 Å spherical probe using the CPPTRAJ surf command (88). To determine the accessible surface area to a larger probe at the residue level, ten hierarchical ensemble 3D structures were generated using all three productions of each system, and then probed for their ability to interact with a 10 Å spherical probe using NACCESS(42, 43, 88). Data are displayed as violin plots, and statistical significance was calculated using either Welch’s t-test or Mann-Whitney U test, depending on distribution normality, using Python. All scripts are available upon request. Figures were generated using GraphPad Prism 9 software.

### Competitive Enzyme-Linked Immunosorbent Assay (ELISA)

To determine the needed concentration of the monoclonal antibody against the Ag85 Complex, a direct ELISA was first conducted. Plates were coated with 2.0 µg/mL of nonglycosylated Ag85B and incubated overnight at 4°C. Plates were blocked with 40 µl/well of 3% bovine serum albumin (BSA) in phosphate-buffered saline (PBS) for 2 hours on an orbital shaker at 100 rpm at room temperature. 20 µl/well of primary antibody (anti-*Mtb* Ag85 Complex IgG1, Clone P4B3G2, BEI Resources) was added using a 2-fold serial dilution and incubated for 2 hours on an orbital shaker at 100 rpm at room temperature. 20 µl/well of a 1:2000 dilution of anti-mouse IgG alkaline phosphatase (AP)- conjugated secondary antibody (Southern Biotech Cat. #1030-04) was added and incubated for 1 hour on an orbital shaker at 100 rpm at room temperature. All samples were run in technical replicates.

After determining the required serum concentration, the competition ELISA was performed. Plates were coated with 2.0 µg/mL of nonglycosylated Ag85B and incubated overnight at 4°C. Plates were blocked with 40 µl/well of 3% bovine serum albumin (BSA) in phosphate-buffered saline (PBS) for 2 hours on an orbital shaker at 100 rpm at room temperature. While blocking was taking place, on a separate plate, competitors (nonglycosylated and glycosylated Ag85B) with four different concentrations of 100 µg/mL, 50 µg/mL, 10 µg/mL, and 1.0 µg/mL were incubated with the determined 1:25600 antibody dilution for 2 hours on an orbital shaker at 100 rpm at room temperature. Afterwards, 20 µl/well of the antibody-competitor solution was added to the blocked plate in technical triplicate and incubated for 2 hours on an orbital shaker at 100 rpm at room temperature. Next, 20 µl/well of a 1:2000-diluted anti-mouse IgG-AP-conjugated secondary antibody (Southern BioTech Cat. #1030-04) was added and incubated for 1 hour on an orbital shaker at 100 rpm at room temperature. Plates were developed using 20 µl/well of 2 mg/mL AP substrate (Sigma Cat. #S0942-200TAB) diluted in 1 M Tris Base, 0.3 mM MgCl_2_ buffer, and OD readings were determined using the BioTek Synergy Neo with the Gen5 Microplate Reader and Image Software. All samples were diluted in PBS-T containing 1% BSA. All plates were washed with 1% BSA in PBS-T between each incubation step. Figures and statistical analysis were completed using GraphPad Prism 9 software.

### Biolayer Interferometry (BLI) for Antibody-Binding Kinetic Assay

BLI assays were performed on a Sartorius Octet R8 instrument at 30°C with shaking at 1000 rpm. All measurements were adjusted by subtracting the background signal from blank loads. Anti-*Mtb* Ag85 Complex IgG1 (Clone P4B3G2) was immobilized on anti-Mouse Fc Capture (AMC) biosensors (Sartorius, Cat. # 18-5088) at 10 µg/ml in kinetic buffer (PBS, 0.02% Tween-20, 0.1% BSA, 0.05% sodium azide). After loading, biosensors were incubated in the kinetic buffer for 10 minutes to establish a baseline signal. Biosensors were then incubated in various concentrations (500, 200, 100, 50, 25, and 10 nM) of the analytes (nonglycosylated Ag85B, glycosylated Ag85B) for kinetic analysis. Curve fitting to determine KD, k_on_, and k_off_ for each analyte was performed using a global 1:1 binding model in the Octet Analysis Software, with R^2^ values exceeding 95% confidence. Biosensors were conditioned before antibody loading and regenerated after the kinetic assay, according to the manufacturer’s recommendations. Figures were generated using GraphPad Prism 9 software.

### PBMC Isolation and Culture

Peripheral blood mononuclear cells (PBMCs) were freshly isolated from healthy donors using SepMate™ tubes (STEMCELL Technologies, Cambridge, MA) according to the manufacturer’s instructions. Briefly, 15 mL of Lymphoprep™ density gradient medium (Catalog # 18060 STEMCELL Technologies) was added through the central insert of each SepMate™ tube. Whole blood, diluted 1:1 with Dulbecco’s PBS (Corning, 21-031-CM), was carefully layered onto the density medium and centrifuged at 1200 × g for 10 min at room temperature with the brake on. Following separation, residual red blood cells were lysed using ACK lysis buffer (Gibco, A1049201).

PBMCs were cultured in RPMI 1640 medium (Corning) supplemented with 2 g/L sodium bicarbonate, 50 μM 2-mercaptoethanol, 2 mM L-glutamine, 1 mM sodium pyruvate, 1% non-essential amino acids, 1% penicillin–streptomycin, and 10% heat-inactivated fetal bovine serum (FBS).

### Cell Proliferation Assay

For proliferation analysis, PBMCs were labeled with 1 μM CellTrace™ Violet (CTV; C34557, Invitrogen) according to the manufacturer’s protocol. Labeled cells were plated at 2 × 10[ cells per well in 200 μL complete RPMI in 96-well U-bottom plates, with each condition set up in triplicate. Cells were stimulated with recombinant Ag85B protein (5 μg/mL) produced either in BL21 bacteria or Expi293F cells. After 72 hours, cultures were supplemented with fresh medium containing IL-7 and IL-15 (5 ng/mL each). Cells were harvested on day 5 for analysis. Proliferation was assessed by flow cytometry (Cytek Northern Lights) based on dilution of CTV within CD4⁺ T cells. Cells were stained with anti-human CD4 antibody (BioLegend), and proliferating cells were defined as CTV_low_ within the CD4⁺ gate.

### Lectin Blotting

Glycosylated Ag85B was treated with α2-3,6,8 neuraminidase (NEB, P0720S) according to the manufacturer’s instructions. Equal amounts of glycosylated and non-glycosylated proteins were separated by SDS-PAGE and transferred onto a membrane. The membrane was blocked in 5% bovine serum albumin (BSA) in Tris-buffered saline with Tween-20 (TBS-T) and incubated with Recombinant Human Siglec-9 Fc Chimera Protein (R&D Systems, CF, Cat# 1139-SL) at a 1:10,000 dilution. After washing with TBS-T, HRP-conjugated human IgG was used as the secondary reagent. Signal was detected using a chemiluminescent substrate.

## Data availability

The mass spectrometry, proteomics, and glycoproteomics data generated in this study will be deposited in a publicly accessible proteomics repository (e.g., PRIDE/ProteomeXchange) upon acceptance of the manuscript, and accession numbers will be provided prior to publication. Molecular dynamics input files, analysis scripts, and processed datasets supporting the findings of this study are available from the corresponding author upon request. All other data supporting the conclusions of this article are contained within the manuscript and its supplementary materials.

## Author contributions

M.S.C. and T.M.A. contributed equally. M.S.C. and T.M.A. designed the study, acquired and analyzed data, interpreted results, and wrote and revised the manuscript. Z.N., E.S.D., M.E.D., and A.P.K. acquired and analyzed data and assisted with interpretation. S.M.N. contributed to study design and acquired and analyzed data. B.R.R. contributed to study design and manuscript revision. A.O. contributed to conceptualization and study design. F.Y.A. conceived and supervised the study, secured funding, contributed to study design and data interpretation, and wrote and revised the manuscript. All authors reviewed the manuscript and approved the final version.

## Supplementary Figures

**Supplementary Figure 1:** A representative spectrum of N52 glycan site demonstrates the rich fragmentation generated from the stepped high-energy collisional dissociation (sHCD), which provides functional fragments both of the peptide backbone and of the glycan itself in a single MS2, which are used by the search engine pGlyco3 to identify glycopeptides.

**Supplementary Figure 2: Ag85B is microheterogeneously glycosylated when expressed in human Expi293 cells.** Site-specific microheterogeneity of **(a)** N52, **(b)** N224, **(c)** N234, and **(d)** N280 using glycomics-informed glycopeptidomics. Relative spectral counts were normalized to the most abundant glycoforms present at each site. Complex structures are represented in dotted orange, hybrid structures are represented in checkered purple, and high-mannose structures are represented in triangle light green.

**Supplementary Figure 3:** Hierarchical clustering of molecular dynamics simulations for the **(a)** nonglycosylated and **(b)** glycosylated Ag85B in ten representative conformations.

