## Supplementary Figures for "Molecular mechanisms of immune evasion by host protein glycosylation of a bacterial immunogen used in nucleic acid vaccines"

### Supplementary Figure 1

293-F Ag85B N52 –H3N4F1 composition MS2

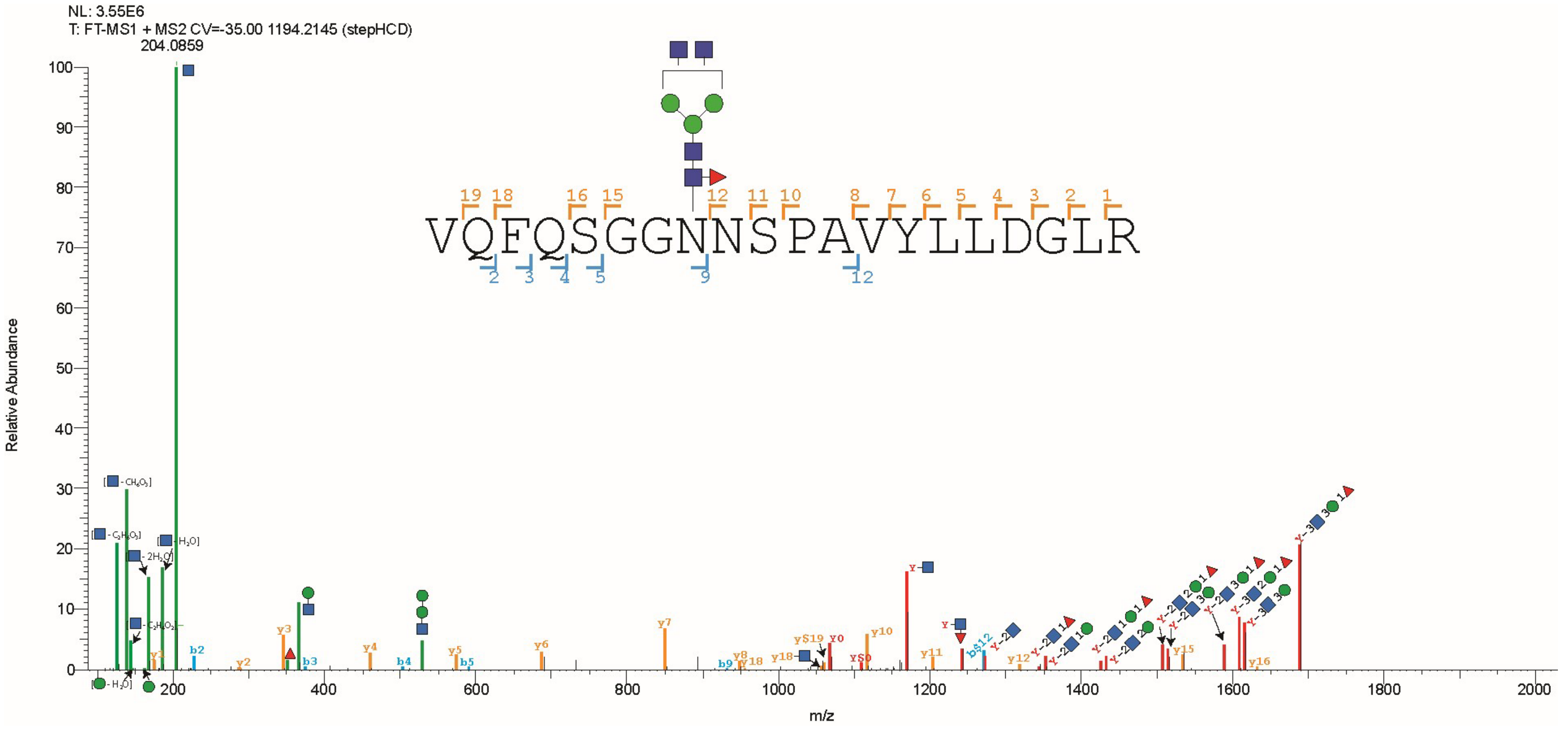

### Supplementary Figure 2

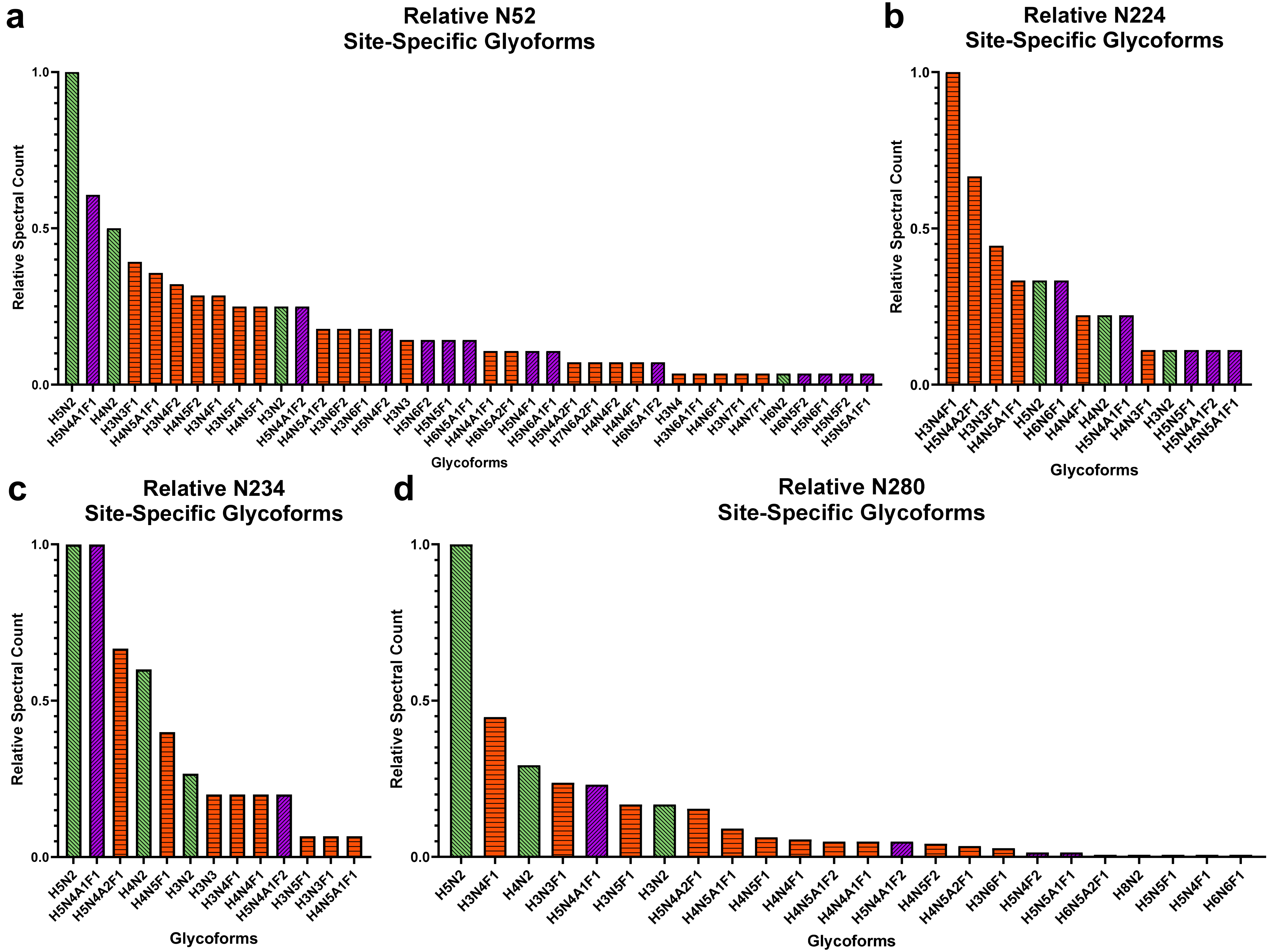

### Supplementary Figure 3

**a** **Nonglycosylated**

**b** **Glycosylated**

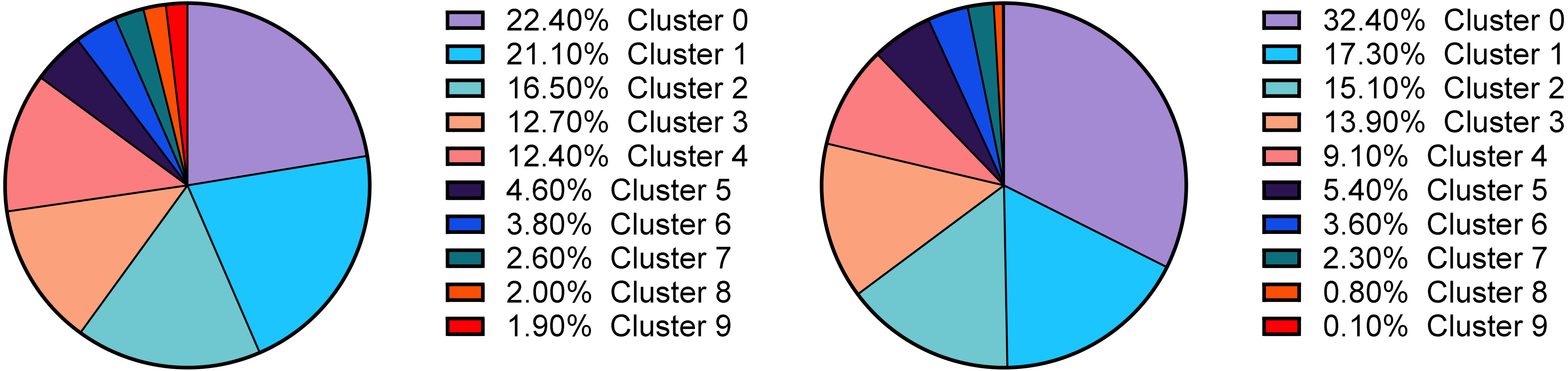
